# Efficient mesoscale phenotypic screening in cultured primary neuron culture

**DOI:** 10.1101/562298

**Authors:** David Nedrud, Yungui He, Daniel Schmidt

## Abstract

A longstanding question in neuroscience is how the activity of ion channels shapes neuronal activity and, as a result, computation in circuits and networks. Optogenetic reagents are tools to answer this question by enabling precise and dynamic perturbation of cellular states. However, development of these reagents can be hampered by low-throughput assays in non-physiological contexts. Here, we develop an all optical phenotypic screen in cultured primary hippocampal neurons that enables the functional assessment of large libraries of genetically encoded optogenetic actuators. Combining real-time analysis and data reduction methods allows for continuous observation of several thousand neurons for several days without onerous data storage overhead. This screening system may be useful in a diversity of research questions that can be coupled to optical perturbation and sensing.

## Introduction

Neurons are excitable cells that form the basic units of biological computation. The molecular basis for their excitability is different types of ion channels and receptors. Changes in activity levels, for example caused by post-translational modification or changes in gene expression, sculpt the function of neural circuits and –ultimately– behavior. In recent years, several opto- and chemogenetic approaches have been developed to change the cellular states of specific types of neurons. Some act through secondary messengers ^1–4^, or by altering gene expression Others consist of exogenous pumps or channels that are expressed heterologously ^9–12^. A third set of approaches directly modulates the activity of endogenous channels ^13–17^. We previously have added a tool called lumitoxin to this toolbox ^18^. Lumitoxins are genetically encoded, membrane tethered peptide toxins that can be actuated with light via an *Avena sativa*(As) LOV2 photoreceptor domain. Illumination causes the C-terminal Jα helix of LOV2 to partially unfold, increasing the flexibility of the tether that connects to the peptide toxin. The result is a decrease in the local concentration of the peptide toxin near the plasma membrane, which ultimately causes the targeted ion channel to become unblocked. We could show that lumitoxins are modular, in the sense that ion channel specificity could be altered by swapping out the encoded peptide toxin. Nevertheless, improvements are required to increase the utility of this tool. Foremost, targeting of ion channel families other than voltage-dependent K^+^channels. Second, an optogenetic reagent that blocks an ion channel after illumination –instead of unblocking them– would be a better fit to commonly used perturbation paradigms in neuroscience. It would also address the concern that lumitoxins, by binding to the targeted ion channel in the resting (dark) state, can affect cellular homeostasis.

Given that the previous approach through which lumitoxins were engineered involved tedious trial-and-error as well as heterologous expression of the targeted ion channels in cell lines, we explored a high-throughput phenotypic screen in neuron culture to streamline this process. We reasoned that cultured neurons are an appropriate system because they express most ion channels and receptors, at physiological levels, that we might want to modulate with lumitoxins. In addition, phenotypic screens that leverage physiological modes of action have had better yields for finding reagents that are effective when compared to target-based approaches involving non-physiological assays ^19^.

One type of phenotypic screening in culture neurons, constellation pharmacology, has been shown to be a valuable approach for discovering new bioactive peptide toxins ^20^. The central idea of constellation pharmacology is that differences in cell-specific channel and receptor expression–collectively called a constellation– create distinct functional phenotypes ^21–24^. It is therefore ion channel diversity itself within populations of cells that provides screen-able content, i.e. distinguishable and characteristic responses after the addition of a pharmacological agent that perturb their function. Functional calcium imaging provides the means to collect this content in a high-throughput manner. However, several technical limitations stand in the way, including the need for manual manipulation (e.g., addition of the pharmacological agent).

Here, we develop an all optical phenotypic screen in cultured primary hippocampal neurons. We introduce real-time analysis and data reduction methods, which allow for continuous observation of several thousand neurons for several days without onerous data storage overhead. Our analysis of calcium imaging data in neurons after the addition of free peptide toxins suggests that toxin effects can be categorized according to ‘fingerprints’. When we train a classification model on these toxin fingerprints, we can detect those neurons –among many transfected with a library of linker and toxin shuffled lumitoxins– that most closely resemble a desired phenotype. In this way we identify a voltage-dependent Na^+^ channel-specific lumitoxin that recapitulates the effects of its free peptide toxin analog in a genetically encoded package. Our phenotypic toxin screening assay sets the stage for linking genotype and phenotype in future studies with more complex library designs. How this assay can be adapted to other research questions that can be coupled to an optical readout is discussed.

## Results

We used cultured hippocampal neurons, which are known to spike spontaneously after several days in culture ^25–27^. Much of their ion channel and receptor complement expression over time has been extensively studied ^28–32^. For an all-optical screen, both sensor and actuator need to be light-sensitive (**Fig. 1A**). Because lumitoxins contain a blue light-sensitive LOV2 domain ^33^, we chose a red-shifted genetically encoded calcium indicator (GECI) RCaMP1.07 ^34^, whose action spectrum does not overlap with AsLOV2 (**Supp. Fig. 1**). Of course, calcium transients are only an indirect product of changes in membrane voltage ^35, 36^, but GECIs have proven useful in many past and contemporary phenotypic screens ^37^.

**Figure 1.**
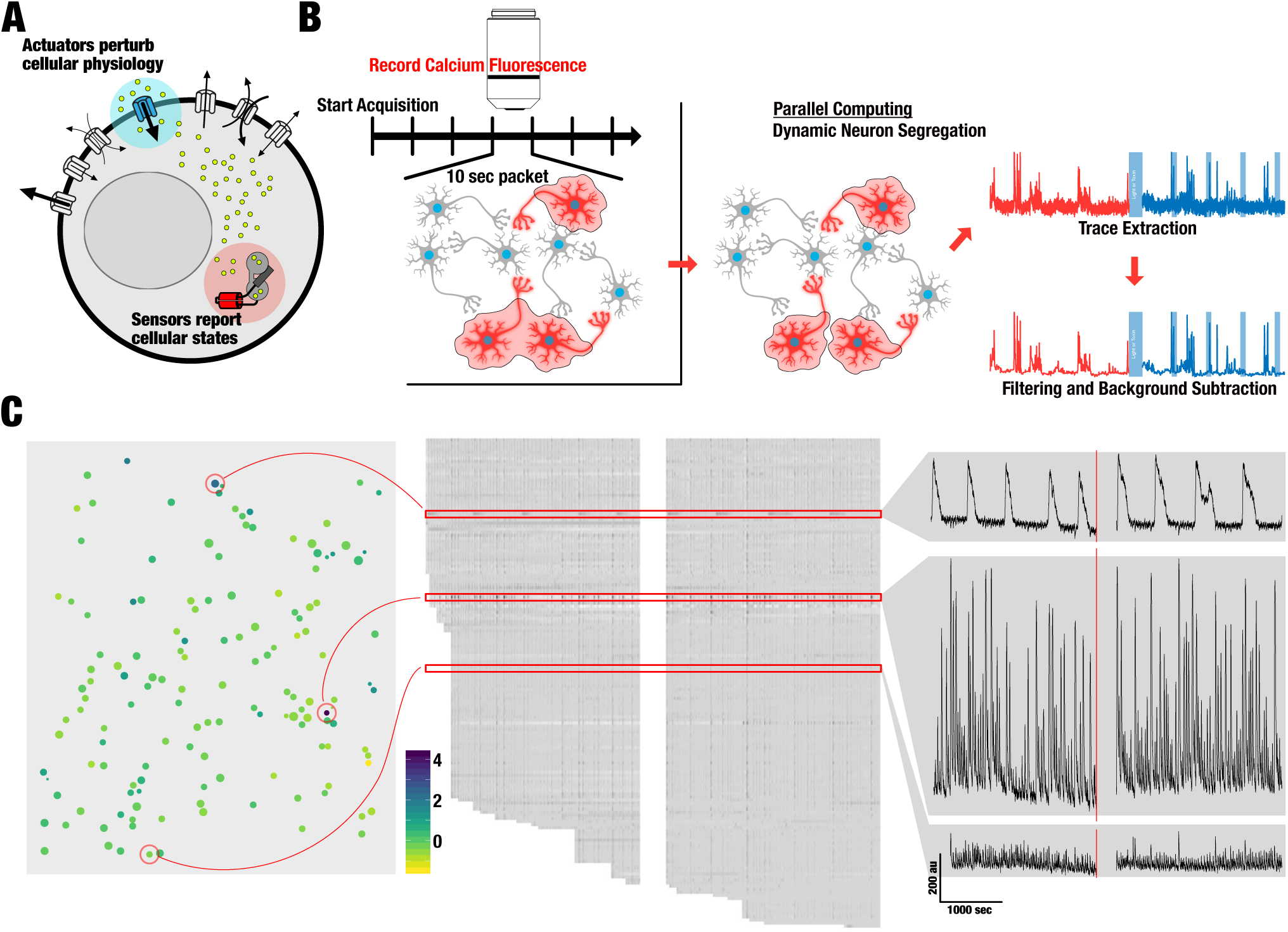
**A. The optogenetic principle.** Cellular physiology is impacted by the constellation of ion channels and receptors (white) expressed on the cell surface that conduct different types of ions and other solutes. Genetically encoded actuators that are controlled by light perturb cellular physiology by providing an exogenous means of conduction across the cell membrane. Genetically encoded optical sensors report changes in cellular states that are caused by this perturbation. **B. Realtime Neuron detection in primary neuron culture.** A live stream of functional calcium imaging data is broken up into packets. Each data packet is segmented into regions of interest (ROI) that represent firing neurons. For ROI that are larger than typical neurons further dynamic thresholding is applied, such that a larger ROI (representing more than one neuron) can be broken down into ROI that represent individual neurons. Trace data for each ROI is stitched together from all packets before (red) and after (blue) a perturbation (e.g. light stimulation, peptide toxin addition). Data is lowpass filtered and stored. **C. A representative reconstructed activity profile for one field of view.** Neurons position within the field of view are indicated (left panel) and color-coded by spike amplitude. Reconstructed traces for all neurons in this field of view (center panel) and detailed views of example traces (indicated by red boxes and circles) are shown (right panel).

Observing GECI fluorescence in parallel for dozen neurons per field is straightforward. All that is required is a motorized microscope system with appropriate climate, illumination, and stage control. We developed custom software based on Matlab and the popular μManager API – available at [link to Github archive forthcoming]– to integrate all microscope components and automate data acquisition. Spike detection, if it were to be done *posthoc* on recorded movies, would require extensive data storage. For example, storing 100 minutes of a 640 by 540 pixels field of view at 10 frames per second requires 100 GB when encoded in the MPEG-1 standard. This problem can be simplified once we realize that most of the recorded field of view does not contain neurons. If we can segment captured data into regions of interest (i.e. individual neurons) in real time, we are able to store only relevant information and discard unneeded pixel information.

We implemented the necessary segmentation of neurons using dynamic thresholding of 100 frames at a time taken from a live camera stream (**Fig. 1B**, **Supp. Fig. 2A**). We quickly realized that standard thresholding methods were not able to segment areas that contained many neurons into distinct single region of interest (ROI). Principle component methods also failed as they required a high amount of computing time that prevented real-time analysis. We therefore developed a simple dynamic thresholding algorithm that employs graduated thresholding and that was able to segment dense neuron populations. The only manual input required is a user-set calcium transient signal amplitude threshold to eliminate false positives. For any ROI that is greater than a standard neuron size (150 pixels at 10x magnification), we dynamically increase the signal threshold for these ROI and continues this loop until all ROI areas are within an expected neuron size. This ensures that neuron-dense regions are properly segmented into individual neurons. The result of this real-time segmentation is a set of ROI for a given data stream excerpt. Only pixel information for these ROI are stored. This loop of segmenting data stream excerpts is repeated as many times as desired by the user to establish a phenotype baseline. In our case this was 30 times for a total of 5 minutes of baseline data. For each loop, neurons are either rediscovered, new neurons are discovered, or neurons that were detected in previous excerpts do not have activity. Pixel information for each of the three categories is recorded. Including the latter category means that any ROI that had activity once, is recorded for the remainder of the experiment, even if the neuron never fires again. Stitching all data excerpts together results in a time series for each detected neuron. **Fig. 1C** shows a representative example of neuron activity in a field of view for 10 minutes, reconstructed from packages of 10 seconds at a time.

After having recorded a field of view for 5 minutes, we introduced a perturbation. This took the form of either manually adding a free peptide toxin to different concentrations (described in more detail below) or illumination with blue light to drive a channelrhodopsin or to switch a genetically encoded lumitoxin. We then recorded for another 5 minutes in the presence of the perturbation. Calcium imaging time series before and after perturbation represent control and treatment datasets, respectively.

We characterized performance characteristic of this data acquisition algorithm (**Supp. Fig. 2B**) and found that data storage scaled favorably, even for very long recordings. Detection for up to 500 objects required less than 5 seconds of computation time. Increasing excerpt length increased computations time, but also increased sensitivity (more neurons detected). The natural ceiling for how many neurons can be detected is a function of magnification and culture density, and we have found that a typical excerpt size of 100 frames (10 seconds at 10 Hz) is adequate.

Next, to detect individual calcium transients in recorded fluorescence data for each ROI, we adapted a template matching algorithm (**Fig. 2A**) ^38^. Briefly, a set of templates is generated from the recorded calcium imaging dataset. Then a sliding window is applied to each trace and compared to this template set. When a correlation threshold is met, the transient found at the current window position is assigned to this spike class. We found that for the signal noise typical in our assay, template matching outperformed other spike sorting techniques such as manual detection, or peak detection by threshold or prominence (**Supp. Fig. 3**).

**Figure 2.**
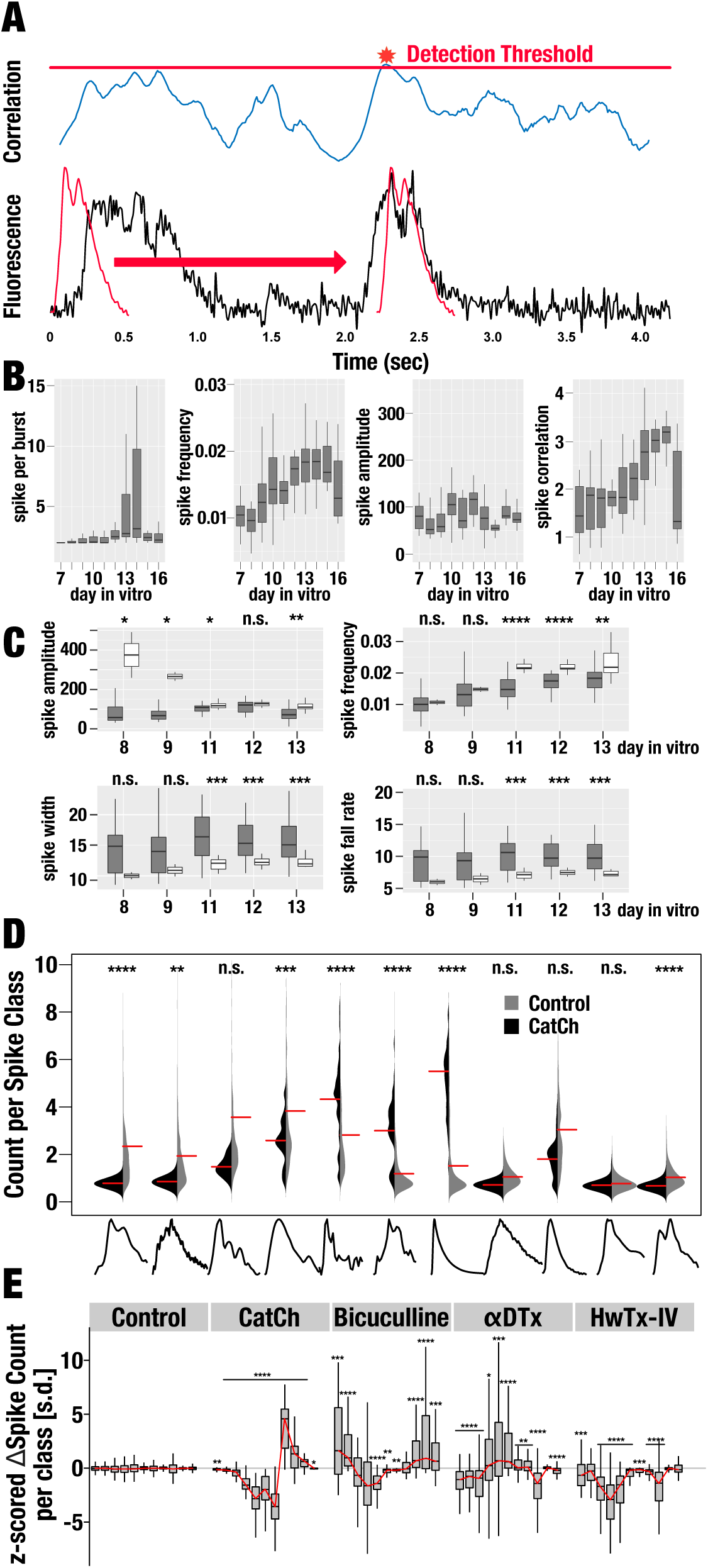
**A. Calcium transient detection by template correlation.** Correlation (top row) is calculated between each calcium transient template (red, bottom row) and each time point in the reconstructed trace (black, bottom row). Correlation coefficients greater than 0.81 are considered a match. If several templates are matched, the one with the highest correlation coefficient is assigned. **B. Developmental changes in primary neuron culture.** Select bulk properties derived from calcium transient of control neurons (i.e. not expressing any exogenous protein) show onset of activity after 8 days in culture, and declining activity after more than 14 days in culture. Box range represents interquartile range, median is indicated by a thick horizontal line, data range is indicated by a thin vertical line. **C. Expressing exogenous proteins alters developmental changes.** Select bulk properties are shown for control neurons (grey) and neurons that express CatCh (white). Significance of the difference between control and CatCh neurons for given day in vitro is indicated at the top of each panel (unpaired Wilcoxon rank sum test, **** p-value < 0.0001, *** p-value < 0.001, ** p-value < 0.01, *, p-value < 0.05, n.s. not significant). **D. Calcium transients in CatCh-expressing neurons.** Beanplots of calcium transients class usage for control neurons (grey) and CatCh-expressing neurons (black). Red horizontal line represents the median. Class template are drawn below and significance (unpaired Wilcoxon rank sum test) of class usage differences are shown above each beanplot. **E. Class utilization fingerprints.** Boxplot of z-scored changes calcium transient class usage for control neurons, neurons expressing CatCh that are exposed to blue light, or neurons subjected to different peptide toxins. Significance of the class utilization difference compared to the same class in control neurons is indicated (unpaired Wilcoxon rank sum test).

One advantage of continuous long-term observation such as we have implemented here is the ability to observe onset and development-dependent changes in network activity. For example, neurons activity is relatively rare until day 9-10 when there is a noticeable uptick in spike frequency and, to a lesser degree, amplitudes (**Fig. 2B**). This is consistent with known expression dynamics of ion channels in cultured neurons ^27, 39, 40^ and changes in excitation/inhibition balance ^41, 42^. Furthermore, synchronized burst become more prevalent after 12 days in cultures, presumably because of continued formation of synaptic connections that can result highly connected networks ^43, 44^. After more than 14 days in culture, spontaneous activity decreases rapidly, which is expected due to accumulation of neurotoxic glutamate in the culture medium (which is not replenished).

Several reports indicated that expressing heterologous proteins, for example Channelrhodopsin (ChR2) ^12^, may change network homeostasis ^45^. To test whether we can observe this in our system, we compared development changes in neuronal activity in neurons expressing the microbial opsin CatCh to those that did not express any heterologous protein. We detected several developmental differences, most notably spike amplitudes, frequency, width, and fall rates (**Fig. 2C**). Spike utilization was also noticeably different, with large amplitude, slow-decaying transient being significantly more often in neurons that express CatCh (unpaired Wilcoxon rank sum test, p-value 2.1 × 10) (**Fig. 2D**). We take these results as an example of retroactive effects that have been described in the context of neuronal excitability; the mere expression of a genetically encoded actuator can alter the baseline behavior of the whole system and must be accounted for in the phenotypic screens ^47, 48^.

In addition to following developmental changes, we wanted to use known modulators of ion channels and measure their effect on network activity. We chose Bicuculline (inhibits GABAA receptors), α-Dendrotoxin (αDTx) (inhibits certain voltage-dependent K^+^ channels), and Huwentoxin IV (HwTxIV) (inhibits certain voltage-dependent Na^+^ channels) all of which are available from commercial sources. Our system robustly detected drastically altered network activity resulting from the addition any of these modulators to cultured neurons. Whereas neurons to which only tyrode was added did not differ in spike class utilization before and after treatment, bicuculline addition increased utilization of many classes consistent with the disinhibiting effect that block of GABA_A_ receptor is predicted to have (**Fig. 2E**). αDTx is predicted to have a similar disinhibiting effect, due the block of Kv1 channels to counteract Na^+^ channel-mediated depolarization. Interestingly, different (compared to Bicuculline) spike classes become more heavily utilized when αDTx was added. Lastly, addition of HwTxIV, which is expected to inhibit activity by virtue of blocking Na_v_ channel, decreases use of all spike classes. When neurons that express CatCh are exposed to blue light –which opens the CatCh pore, allows cations to flow across the cell membrane, and depolarizes the cell– utilization of large amplitude, slow-decaying calcium transient was increased at the cost of all other classes that were used in CatCh-expressing neurons in the absence of illumination. Altogether, from changes to spike class utilization alone, we can ascribe ‘fingerprints’ to each perturbation (GABA_A_ block, K_v_1 block, Na_v_ block, or exogenous cation ion channels opening).

To further test if we could measure modulator-specific effects on bulk firing properties, we further parameterized each detected calcium transient (amplitude, fall & rise time, etc.) for each spike (**Fig. 3A**). We also measured how correlated each neuron’s activity was with the rest of the network using the mutual information statistic ^49^ (**Supp. Fig. 4**) and cross correlation. All bulk firing property data is time-averaged for each condition (control and treatment) associated with identifiers for neurons, day in vitro, field of view, coordinates, biological replicate. In total, we measured 10 properties that describe activity before and after perturbation (light stimulation or peptide toxin addition). These properties are: amplitude, frequency, width at half height, rise rate, fall rate, burst frequency, events per burst, Fourier power distribution, and network properties - mutual information, correlation. While we could discern gross trends from manual inspection of this data –for example, HwTxIV addition decreased overall spontaneous firing and αDTx drastically decreased firing frequency regularity– we suspected that much of the useful information was spread across several measured variables. We therefore subjected this dataset to principal component analysis to reduce dimensionality. Much of the variation (75%) on the dataset is explained by just the first three principal components (**Fig. 3B**), allowing us to cluster response to perturbation along two or three dimensions (**Fig. 3C**). Using PCA scores to select neurons that score different than 95% of the control neurons, we could isolate from hundreds of neurons those are the strongest responders to a perturbation (**Fig. 3D**).

**Figure 3.**
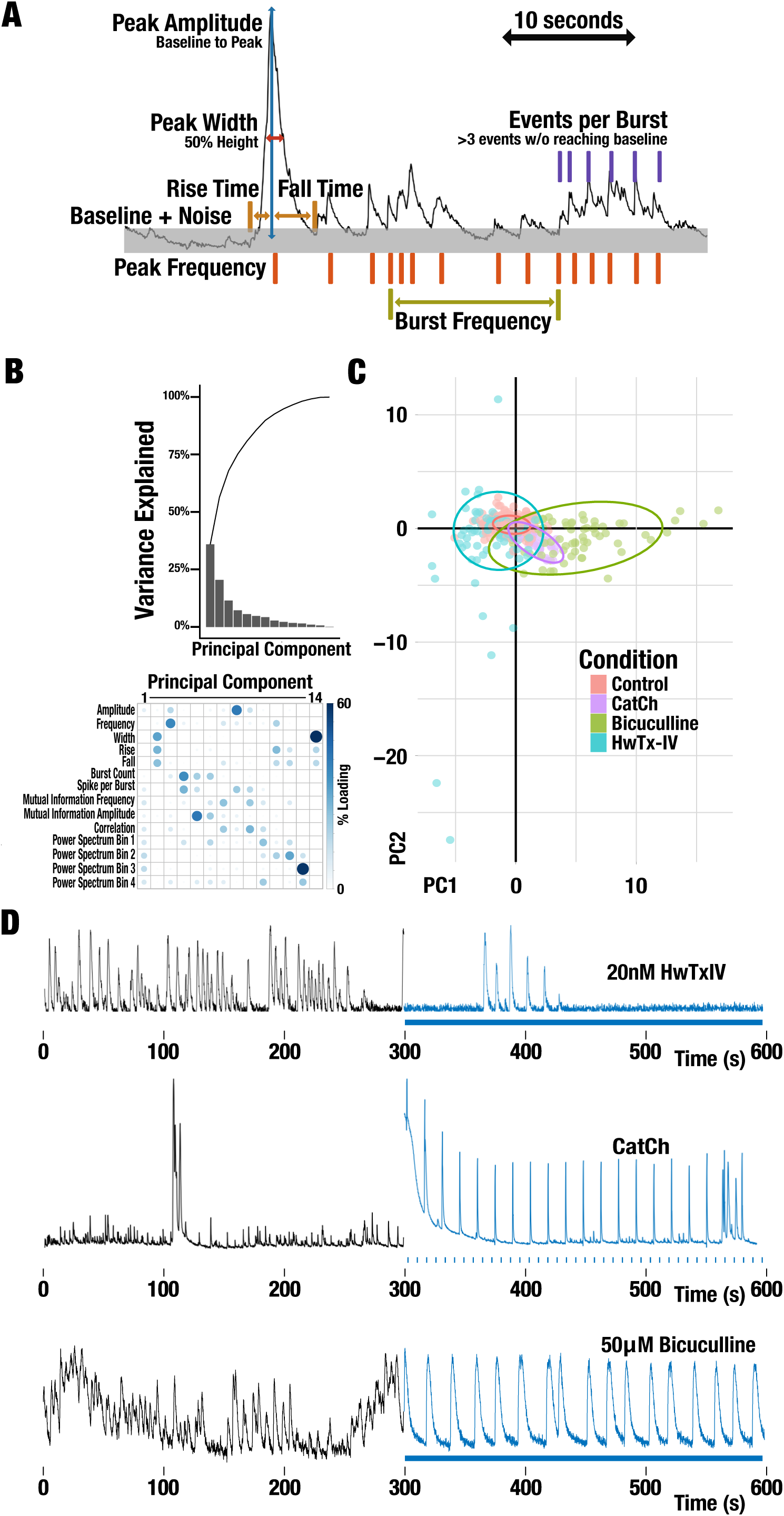
**A. Calcium transient parametrization.** Each detected calcium transient is further analyzed to extract several characteristic measures (indicated in bold font). **B. Principle Component Analysis of calcium transient parameters.** Scree plot of explained variance (left panel) and loadings for each principle component (right panel). **C. PCA allows clustering of neurons subjected to different type of perturbation.** PC1 and PC2 are plotted for control neurons (red dots), neurons expressing CatCh stimulated with blue light (ochre dots), and neurons subject to different peptide toxins (inhibiting GABAA receptors, green dots; inhibiting Kv1.x channels, blue dots; inhibiting Nav1.x channels, purple dots). Normal confidence ellipses (confidence level 0.95) for each condition are shown (shaded). **D. Representative Traces before and after perturbation.** Functional calcium imaging traces are shown when 20 nM HwTxIV is added (top), 50 μM Bicuculline is added (center), or a CatCh-expressing neuron is stimulated with blue light. Black and blue trace portions indicated RCaMP1.07 fluorescence before and after perturbation, respectively. For CatCh, light was pulse at 0.05 Hz with a 2.5% duty cycle.

Since the effect of a perturbation (light stimulation or peptide toxin) could be clustered into distinct groups, we further probed whether we can train a classification model that would be useful in predicting the type of toxin that is causing a specific change in firing properties. We chose decision tree classification models, which are good at capturing non-linear interactions between descriptive variables. Consistent with the first three principal components explaining most of the variance, decision tree models predominantly used these components in classifying changes in neuron firing properties based on how this neuron was perturbed (**Supp. Fig. 5**). Receiver operating characteristics (ROC) and confusion matrix testing show that model performance is acceptable (Multi-class area under the curve: 94.21%, Accuracy 94%).

Now that we could predict the type of perturbation that had caused a set of changes in bulk firing properties of neuron, we tested whether this system would be useful for a phenotypic screen of new kind of lumitoxins. To this end we synthesized lumitoxins that genetically encoded 84 different peptide toxins spanning different sources (spiders, scorpions, etc.) and target different channel families (K^+^, Ca^2+^, Na^+^, etc.). We also varied the linker through which a toxin is tethered to the membrane-bound LOV2 domain. Total library complexity was 17 linkers × 84 toxins = 1428 variants.

Since peptide toxins contain several disulfide bridges, are often very short (<100 amino acids), and rich in hydrophobic residues, it can be hard for them to achieve their native fold when expressed heterologously, and may not traffic to the surface at all when part of a lumitoxin. We therefore assessed which library members express to the cell surface. After transfection into HEK293 cells, a FLAG tag inserted in between the toxin and the LOV2 allowed us to fluorescently label cells that surface express a lumitoxin via anti-FLAG Alexa 568 antibodies. GFP, expressed on the lumitoxin’s C-terminus served as a transfection marker. By nature of transfection of culture cells with cationic polymers, each cell likely is transfected with several library members (some of whom might surface traffic, while others do not). We therefore employed a quantitative NGS approach to connect genotype (linker and toxin) to phenotype (surface expression). We sorted cells into Alexa658^high^/GFP^high^ and Alexa568^low^/GFP^high^ populations, from which we isolated plasmid DNA for library preparation and Illumina MiSeq (**Supp. Fig. 6**). We analyzed reads from both populations (normalized to a transfection control) to identify library members (linker and toxin combinations) that are enriched in the Alexa568^high^ population and depleted in the Alexa568^low^ population. The two top hits were α-Dendrotoxin (as expected from previous work ^18^) and HwTxIV (**Fig. 4A**). These data suggested HwTxIV as a good candidate for implementing Na_v_-directed lumitoxins.

**Figure 4.**
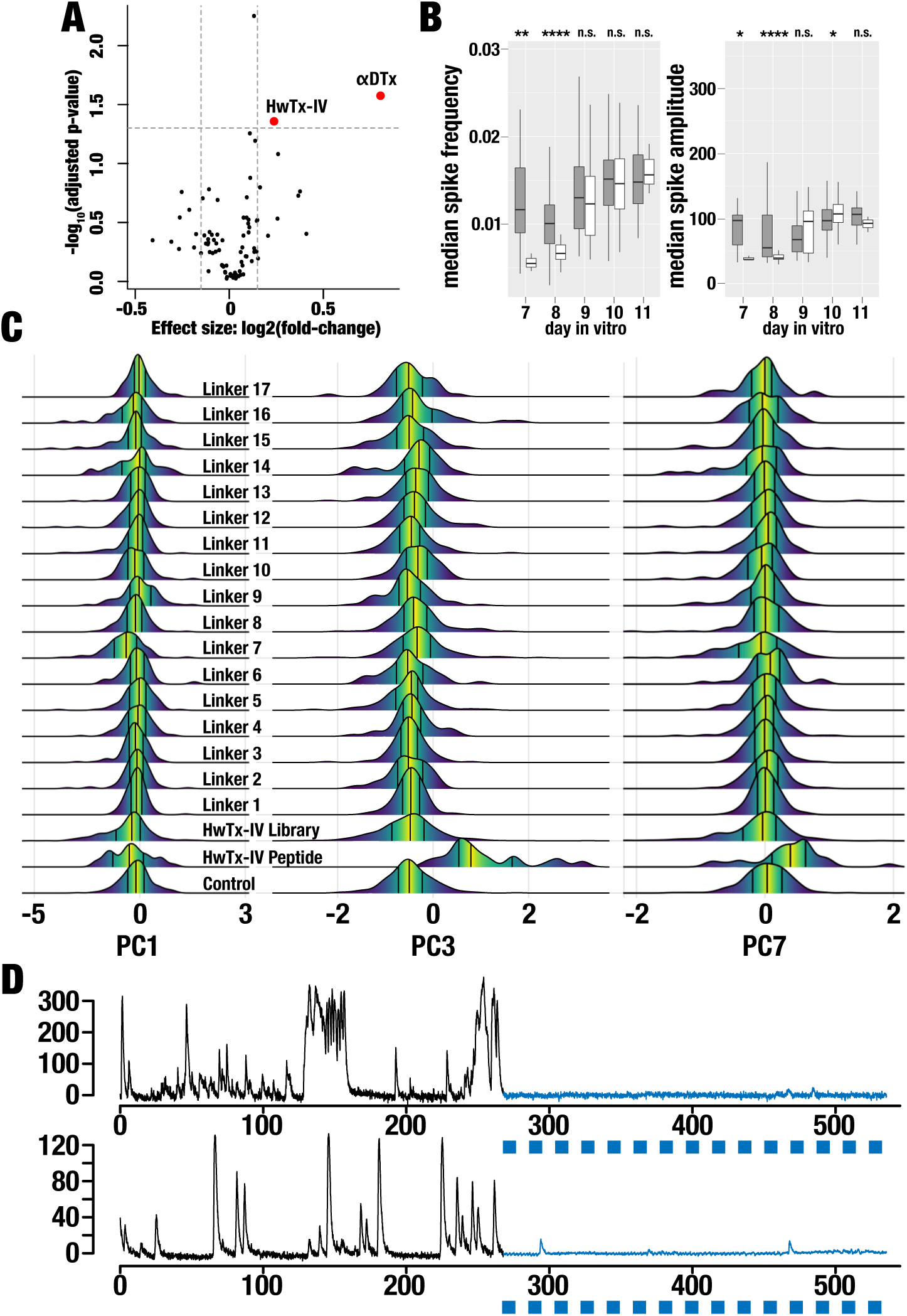
**Peptide toxin that are surface-expressed when encoded as lumitoxins.** Volcano plot of surface expression testing for lumitoxins libraries (represented as dots) that encode one of 84 different peptide toxins and any of 17 linkers. Only libraries that encoded HwTxIV or αDTX where significantly surface-expressed (red dots). **B. Expressing of HwTx-lumitoxin alter onset of developmental changes, but not steady-state bulk properties.** Select bulk properties are shown control neurons (grey) and neurons expressing the complete HwTx-lumitoxin library (white). Onset of spiking is delayed for the latter. Significance of the difference between control and HwTx-lumitoxin neurons for given day in vitro is indicated at the top of each panel (unpaired Wilcoxon rank sum test, **** p-value < 0.0001, *** p-value < 0.001, ** p-value < 0.01, *, p-value < 0.05, n.s. not significant). **C. HwTx-Lumitoxin efficacy is related to linker type.** Ridgeline plots for the top three principle components used in the perturbation type classification tree model. Neurons expressed either a blank lumitoxin (i.e. does not contain a toxin; control), where perturbed with free 20 nM HwTxIV peptide, the complete HwTx-lumitoxin library, or a HwTx-lumitoxin with a specific linker type (Linker 1 – Linker 17, see **Supp. Table 1**). **D. Representative examples of HwTx-Linker7 lumitoxin efficacy.** Functional calcium imaging traces are shown for two neurons infected with HwTx-Linker7 lumitoxin that the perturbation type classification tree model indicated as responding akin to free HwTxIV peptide toxin. Black and blue trace portions indicated RCaMP1.07 fluorescence before and after blue light stimulation (0.05 Hz with a 2.5% duty cycle).

In earlier work we had shown that linker optimization is a critical part of a lumitoxin design ^18^. We therefore asked whether our phenotypic screen would be useful for identifying HwTx-linker combinations that are more effective than others. Furthermore, we wondered whether the same screen could help us identify linker combinations that give lumitoxin a desired ‘blocking with light’ phenotype. We transduced cultured hippocampal neurons with a library of HwTxIV lumitoxin that included several linker types (**Supp. Table 1**). As we had done for free peptide toxins and CatCh, we followed onset of spontaneous activity and characterized baseline development changes. We found that typical developmental changes are delayed (by about 2 days) when HwTxIV-lumitoxins are expressed, but nevertheless spike amplitude and frequency are comparable to controls after 10 days in culture (**Fig. 4B**).

We then measured the effect of blue light illumination (500µW/mm^2^ for 10 seconds followed by pulsed illumination at 0.05 Hz with a 2.5% duty cycle) on neuronal activity. It was clear that a subset of neurons transduced with the lumitoxin library responded with changes in activity upon illumination (**Fig. 4C**). We did not observe this in neurons that express a mock lumitoxin (only LOV2-PDGFR lacking an encoded peptide toxin). We also observed similar changes in some, but not all linkers we tried (e.g., compare linker 7 vs. linker 1 (**Fig. 4C**). Using Dunnetts many-to-one comparison of the top three principle components used in the classification tree (PC1, PC3, and PC7) with mock lumitoxin as a control, light-induced changes were significant for neurons expressing the complete HwTxIV library (p-value 4.3 × 10^−12^), and those expressing HwTxIV-Linker7 (p-value 6.4 × 10^−6^). Two conclusions can be drawn from this. First, HwTxIV appears to be effective at modulating neuronal electrophysiology when expressed as part of a lumitoxins. Second, these light dependent changes appeared to be linked to specific linker used in the construction of lumitoxins, which implies that efficacy of the HwTxIV lumitoxin dependent on a specific linker.

We next used the classification model trained on peptide toxin and applied it to lumitoxin data. Interestingly, 8.2% of lumitoxin library neurons were predicted as HwTxIV-like (**Supp. Table 3**). Less than 1% were predicted as Bicuculline or Channelrhodopsin. As one would expect, the percentage of HwTxIV-like neurons increased for linker in which observed light-dependent activity changes (e.g., Linker 7: 11.2%) but not others (e.g., Linker 2: 1.2%). When we selected neurons that have HwTxIV-like properties and compared compared bulk properties, we found that for Linker 7 light-dependent changes, specifically firing frequency and mutual information, resemble those observed for added free peptide toxin (changes similarly for both free and HwTxIV lumitoxin (**Supp. Fig. 7**). A manual inspection of trace data supported the idea that the classification model could identify, based on bulk properties, neurons that express a specific lumitoxin which is modulated with light (**Fig. 4D**). Similar to what is observed after the addition of free peptide toxin, neurons expressing HwTxIV-lumitoxin showed markedly less firing after onset of illumination.

## Discussion

Expression and activity levels of different ion channels and receptors differ among types of neurons and define their function identity. In recent years a technology framework has emerged that combines optical or pharmacological tools with cell-type specific gene expression. Opto- and chemogenetic reagents enable the interrogation of circuits, and in principle allow us to determine quantitative aspects of ion channels’ contributions to circuit function that will improve mechanistic models of neuronal tissue. Most of these reagents, however, act at the circuit level and there is a particular paucity of tools that can modulate the activity of endogenous ion channels. We previously had developed lumitoxins as a prototype technology to overcome this barrier. It became clear, however, that a more generalizable protein engineering approach was required to diversify this reagent to other ion channels and to improve its functional characteristic. In particular, lumitoxin efficacy in cultured cells lines overexpressing a target ion channel differed markedly from their efficacy in primary cultured neurons where ion channels are expressed at physiological levels. We therefore set out to develop a high-content phenotypic assay, which more closely resemble the physiology of the tissue that lumitoxin will be applied in. We hypothesized that perturbation to any part of the constellation of ion channels expressed in a given neurons will alter its functional output (i.e. spontaneous activity). Accordingly, we developed an all-optical functional calcium imaging assay that allows us to observe neuronal activity and perturb individual types of ion channels either through direct peptide toxin addition or light-switched lumitoxins. We find that our assay scales to thousands of neurons and enables long-term observation (>7 days), while keeping data storage requirements to a minimum. This is because of an efficient region-of-interest focused data acquisition algorithm that extracts and stores only pixel information belonging to active neurons. Spike detection and sorting are often performed post-acquisition because this allows for the greatest flexibility in terms of optimization of the underlying algorithms to maximize sensitivity and precision. Real-time methods have become practical in recent years, in particular when experimental conditions can be simplified (e.g. fixed position in vivo recording, worm tracking, or as described here optogenetic stimulation and cell culture plate scanning).

We observed stereotypical developmental changes in neuronal activity, which were altered when exogenous proteins are expressed. This opens up the possibility to use this kind of long-term all-optical observation as an assay that tests whether the expression of a heterologous protein alters baseline cellular physiology and homeostasis. We could establish the same kind of fingerprints for neuronal activity changes when we added free peptide toxin that block different types of ion channels. When we then trained machine learning algorithms on the changes to bulk neuronal activity properties, we could predict which toxin had caused what type of effect. Together this suggests that with sufficient training data (changes in bulk neuron activity properties in response to commonly used peptide toxins) a generalized model might be established that could be useful for the discovery of novel peptide toxins and related agents. A similar decomposition of zebra fish phenotypic activity data into fingerprints allowed the categorization of small molecules psychoactive drugs based on what type of ion channel or receptor it modulated ^50^. It also enabled the discovery of new entities based on fingerprint similarity ^51^.

We also showed that our approach is useful to diversify the class of lumitoxins to ion channels other than voltage-dependent K^+^ channels. An unbiased screen for folding and surface expression among 84 candidates identified HwTxIV as a candidate. HwTxIV belongs to the inhibitor cystine knot family, which has many biotechnology applications including treatment of neuropathic pain ^52^, imaging and treatment of cancer ^53, 54^, and as growth factor mimetics ^55, 56^. Our data from linker libraries of HwTxIV-Lumitoxin and individual linker variants show linker optimization is required for maximum lumitoxin efficacy. Interestingly, we could identify at least one specific linker, a polyproline motif, that gave the HwTxIV-lumitoxin a ‘block-with-light’ phenotype. That is, with onset of illumination, neuronal activity decreased, just as was observed when free peptide toxin was added. Based on a computational model that describes the light-induced unfolding of the LOV2 Jα as an event that increase the volume a tether toxin can explore, it was suggested that the first generation of lumitoxins acts by decreasing a toxin local concentration close the membrane-embedded ion channel ^18^. We speculate that the rigid secondary structure a polyproline helix might mean that in the dark the toxin is captured in a conformation that is much less competent to bind the ion channel, perhaps keeping the toxin pointed away from the cell membrane. Upon illumination, the unfolding of the Jα helix significantly increases the degrees of freedom the tethered toxin has, which now can bind and block its cognate receptor, voltage-dependent Na^+^ channels. Of course, further biophysical and electrophysiological characterization is required to fully understand the structural basis for this apparent sign switch in lumitoxin function.

## Acknowledgements

We thank the Schmidt Lab for feedback and advice, the University of Minnesota flow cytometry resource and genomics center for assistance. D.N acknowledges funding through the Minnesota Discovery, Research, and InnoVation Economy (MnDRIVE) Fellowship.

## Author Contributions

D.N. and D.S. designed the study. D.N. conducted the experiments. Y.H. provided research support. D.N. and D.S. analyzed the data and authored the manuscript.

## SUPPLEMENTARY MATERIALS

### Materials & Methods

#### Primary Neuron Culture

All animal procedures were in accordance with the National Institute of Health Guide for the care and use of laboratory animals and approved by the University of Minnesota Institutional Animal Care and Use Committee (Protocol #1503-32420A). Hippocampal regions from CD-1-022 mice (Charles River Laboratory) mice postnatal day 0 − 1 were isolated and digested with papain (100 units in Hanks balanced salt solution supplemented with 35mM glucose, 1mM Kynurenic acid, 0.3 mg/ml L-Cysteine and 10mM MgCl_2_) for 6 − 8 minutes. Cell suspension was washed with Ovomucoid trypsin inhibitor (10mg/mL), washed three times with 1 mL of plating media (MEM, 10% fetal bovine serum, 0.5% glucose, 10mM HEPES, 2mM L-glutamine, 0.5mg/ml holo-transferrin, 25μg/ml insulin, B27 supplement, buffered to pH 7.4 with NaOH). The tissue was then mechanically dissociated by triturating through P1000 plastic pipette tips, and settled by gravitation. Dissociated neurons in the supernatant were plated on matrigel coated 24 well glass bottom plates at approximately 50,000 cells per well and maintained in plating medium.

#### Lumitoxin Library Preparation

Cassette encoding for 17 peptide linkers (**Supp. Table 1**) and 84 peptide toxins (**Supp. Table 2**) were synthesized (Genscript). Each cassette is flanked by BsaI restriction sites such that lumitoxins can be assembled from digested PCR products (toxin, linker, lumitoxin backbone) using Golden Gate Assembly ^57^ (**Supp Fig. 8**). An all-toxin/all-linker library (1,428 variants) for surface expression testing was assembled by mixing all toxin and all linkers at equimolar ratio. A HwTxIV-lumitoxin library (17 variants) was assembled by mixing the HwTxIV cassette with all linkers at equimolar ration. Individual HwTxIV-linker variants were assembled from the HwTxIV and a specific linker cassette at equimolar ratio. Individual linker constructs were sequence verified; a random sample was drawn from libraries to verify sequence. Assembled libraries were subcloned into a viral payload shuttle vector using BamHI and EcoRI sites.

#### Virus Production

AAV-DJ ^58^ for delivering the various lumitoxin payloads was packaged ^59^. Briefly, 5.2 × 10^6^ AAV293 cells were triple-transfected 2µg of pAAV-DJ, 3µg pHelper, and 1.7µg of viral payload shuttle vector (encoding lumitoxin or CatCh ^46^ flanked ITR sites). After 72 hours, viral particles were released from producer cells by repeated freeze/thaw cycles in the presence of Benzonase (100 units). Crude lysates were cleared by centrifugation. Viral particles in the supernatant were titered using qPCR with ITR-specific primers. Supernatants were stored at 4degC and used for viral transduction without further purification.

#### Viral Transduction

For all experiments, cells were infected (∼20,000 vector genomes per cell) with AAV delivering RCaMP1.07 driven from the CamKII promoter (UPenn Virus Core). Lumitoxin or CatCh virus was also added at the same time (5,000 − 100,000 vector genomes per cell). Plating media was removed and neurons were washed with MEM twice. Virus was added in 200 µl MEM. Following a one-hour incubation, virus was removed and original media was replaced along with 2 µM AraC.

#### Surface expression assay

HEK293FT cells were maintained in DMEM, 10% fetal bovine serum, 1% penicillin/streptomycin and 1% sodium pyruvate. The all-toxin/all-linker lumitoxin library (100ng) was transfected into 5.5 × 10^5^ HEK FT 293 cells using TurboFect. After 48 hours, cells were detached using Accutase and washed with FACS buffer (2% FBS, 0.1% NaN3, 1xPBS). Cells were then incubated for 1 hour with mouse anti-FLAG antibody at 1:200 dilution in FACs buffer rocking at 4degC. Cells were washed twice with FACS buffer and then incubated with goat anti-mouse Alexa568 antibody at 1:400 dilution for 30 mins rocking at 4degC and protected from light. Cells were again washed twice with FACS buffer and filtered using a cell strainer. Cells were sorted into GFP^high^ / Alexa568^low^ (transfected cells without lumitoxin surface expression) and GFP^high^/Alexa568^high^ (cells with lumitoxin surface expression) on a BD Bioscience FACSAria II flow cytometer (**Supp. Fig. 6**). GFP fluorescence was excited using a 488 nm laser, recorded with a 525±50 nm bandpass filter and a 505 nm long pass filter. Alexa fluorophore 568 fluorescence was excited using a 561 nm laser and recorded with a 610±20 nm bandpass filter. Cells were gated on Side Scattering and Forward Scattering to separate out whole HEK293FT cells, gated on forward scattering area and width to separate single cells, then gated on coexpressed GFP to gate out cells that received a plasmid, then gated on cells that were labeled using the anti-flag antibody for surface expressed channels. Gates were determined using single wildtype, GFP only and unstained library samples. GFP^high^ / Label^low^ and GFP^high^ / Label^high^ cells were collected into catch buffer (20% of FBS, 0.1% NaN3, 1xPBS). Between 9,000 and 60,000 cells were collected for each condition, which represents 7-50 depth of coverage. DNA was recovered from each sample using Quick-DNA microprep kit. Chromosomal DNA was removed by digestion with Plasmid-Safe ATP-dependent DNAse and plasmid DNA was further concentrated using DNA purification kits. Lumitoxin amplicons were prepared using 20 cycles of PCR with Primestar GXL and gel purified. Purified amplicon DNA was quantified using picogreen DNA concentration at the University of Minnesota Genomics Core. Libraries were generated at University of Minnesota Genomics Core using Nextera XT library generation workflows to fragment and add on Illumina sequencing adaptors, and sequenced using MiSeq. On average 200,000 reads per were recorded sample (transfection control, not-surface expressed, surface-expressed) for each of the two biological replicates. Reads were analyzed for toxin and linker identity using pairwise alignment implement with the Needleman-Wunsch algorithm in Matlab. Count tables of toxin and linker pairs were further analyzed using the DESeq2 package ^60^ in R ^61^. Toxins with a p-value less than 0.05 and a 1.2-fold increase over control were considered significantly present on the cell membrane.

#### Functional Calcium Imaging: Hardware

Our data acquisition system is based on a Nikon Eclipse Ti inverted microscope with a 10x/0.45 Nikon CFI Plan Apo objective, an Andor Zyla 5.5 sCMOS camera, a Ludl BioPoint2 stage, a Lumencor SPECTRA X light engine, and an Okolab Boldline stage mounted incubator. The system is house in a custom-built light-blocking enclosure and vibration-isolated. All data acquisition hard is integration through the micro-manager API and controlled by custom code implemented in Matlab.

#### Function Calcium Imaging and Data Acquisition

We have developed custom software for both acquisition and analysis of calcium fluorescence imaging implemented in MATLAB. The software is available on GitHub (will be available at the point of publication). Our workflow begins by loading the micro-manager core java API and a configuration file with user specific hardware. The software then requires user input for acquisition parameters: blue light pulse interval and duty cycle, camera settings (exposure, binning), loop count and duration, cell detection threshold, and number of field of views (FOV). Pulse information is required for blue light stimulation during the repeat acquisition. Camera settings are optimized for calcium/voltage sensor and interval time defines the “real-time” resolution that is repeated for the number of loops. The user-set threshold used for removing false positives is set in the user interface. FOV positions are either automatically selected based on a spiral scan or manually selected with a user interface. Upon acquisition initiation, the software starts a video acquisition in a parallel thread and batches buffered video frames every 10 seconds (interval-time parameter). After the video batch is delivered to the main thread, video is filtered for noise using a 3×3 two-dimensional convolution pixel block averaging for each frame. At this point, a temporary background trace is determined by averaging all pixels excluding previously identified neurons to detect and remove bleed through from the blue light stimulation (see light stimulation). To identify neurons, we detected regions of interest (ROI) based on a dynamic threshold of neuron activity. Calcium activity was identified by calculating standard deviation of each pixel across all frames. Light stimulation can cause increased standard deviation and false positive neuron detection. Therefore, frames with excessive background fluorescence are subtracted from thresholding. Using a user-specified initial threshold, ROIs are formed and area is calculated. ROIs that exceed the area of a standard neuron soma are selected and looped through an iteratively increasing threshold until all somas are within the standardized area range. Active neurons are then aligned with previously detected neurons to identify newly detected neurons. Finally, a calcium trace is extracted from each neuron by averaging all pixels within a ROI and the background fluorescence is recorded from all pixels excluding neuron ROIs. This batching process continues for the duration of the experiment. Raw data files are saved with calcium traces and neuron descriptions.

#### Light Stimulation

Light stimulation was implemented as 500µW/mm^2^ for 10 seconds followed by pulsed illumination at 0.05 Hz with a 2.5% duty cycle. Wavelength was set by a 470±24 nm bandpass excitation filter in the Lumencor light engine.

#### Perturbation with Peptide Toxins or Channelrhodopsin

Peptide Toxins were ordered from Alomone labs (STH-101, D-350, B-136; HwTxIV, αDTx, Bicucculine). For peptide toxin characterization, media was adjusted to 1 mL before adding toxin to account for loss of volume during incubation. Neurons traces were then acquired using the previously described methods. Following a five-minute acquisition, the peptide toxin was added in a sterile hood. Upon returning the plate to the stage, FOV alignment was manually performed by neuron image overlay and optimized with image registration using mutual information and gradient decent. Finally, neurons were recorded for a second five-minute acquisition.

The channelrhodopsin CatCh ^46^, was expressed from a human synapsin promotor. Acquisition of CatCh followed the same protocol as the Lumitoxin acquisition.

#### Functional Calcium Imaging Analysis

For calcium trace analysis we used High Performance Computing infrastructure available at the Minnesota Supercomputing Institute (typically 8 nodes built on Intel Haswell E5-2680v3 processors). Raw data files from the acquisition were imported, grouped by name, and concatenated. During this import, background calcium trace was subtracted from individual neuron traces to account for FOV fluctuations in fluorescence. Calcium transients were detected by template correlation, which was adapted from Patel et al. ^38^. Templates were taken from FluoroSNNAP GitHub and hand-picked from recorded calcium traces from a variety of neuron states. The correlation was calculated between each template for each time point in the trace plus the length of the template (**Fig. 2A**). Correlations that reach above 0.81 are considered a transient and the best matched template is recorded. We parametrized each transient further by frequency, amplitude, width, rise-time, fall-time, Fourier Transform power distribution, and mutual information, and correlation. Frequency [Hz] was calculated from the inter-event-interval. Amplitude (Δ fluorescence [AU]) was calculated as difference between peak height and baseline. Peak-width [seconds] was calculated at half peak height. Rise-time and fall-time [seconds] were calculated from half height to peak and from peak to half height. Fourier transform power spectrum [W/Hz] was used to determine densities of frequency. The spectrum was broken into 4 parts to easily compare distribution of frequencies. Finally, mutual information and correlation provided a parameter by which synchronicity and functional connectivity can be measured. Mutual information and correlation were calculated with pairwise comparisons of raw fluorescent traces for each neuron permutation within a field of view.

This analysis was repeated for control and treatment datasets and each parameter was returned as an average and standard deviation for each neuron. Perturbation-dependent changes were calculated as the difference between treatment and control data for the same neuron. Due to the variability of neuron culture, measurements were normalized to an experimental control within the same plate and time period. For each parameter and each neuron data was normalized by z-scoring:

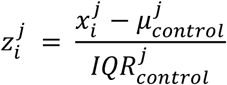

Where *x* is the data for the *J*th parameter and *ith* neuron, *μ* is the mean for the *J*th parameter in control neurons, and *IQR* is the control neuron interquartile range for the *J*th parameter. This resulted in measurement for each neuron being reported as a standard deviation from the control mean.

#### Classification and Prediction

Predictive models were implemented as decision tree using the rpart package in R ^61^. Tree depth was limited 4 levels to minimize overfitting and cross-validated 10 times. PCA-transformed Functional calcium imaging data from neurons to which different peptide toxin have been added, or which expressed CatCh and were stimulated with blue light served as the training dataset. Model performance was assessed by multiclass receiver operating characteristic (ROC) and confusion matrices.

**Supplementary Figure 1:**
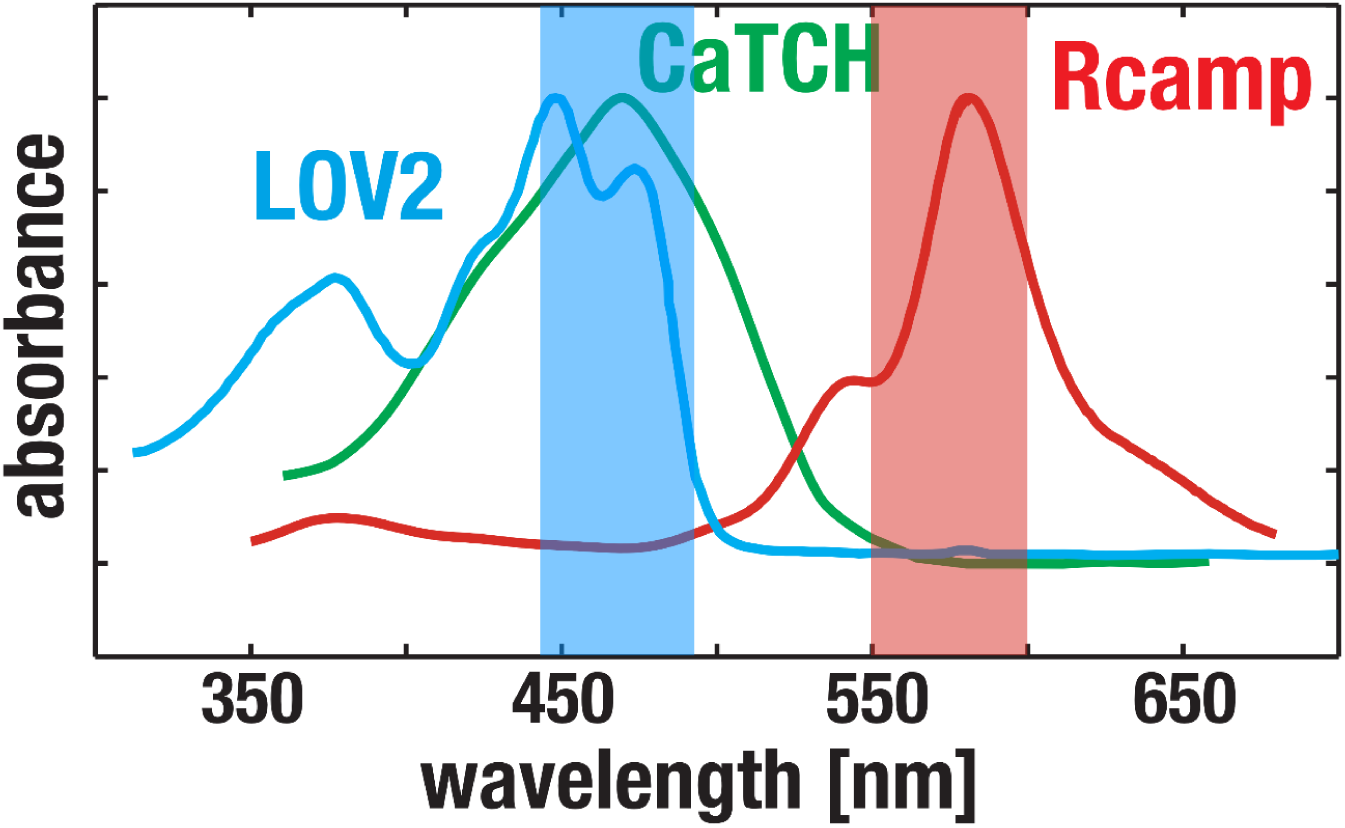
Absorbance spectrum for optogenetic reagents used in this study. RCaMP1.07 was stimulated at 575±25 nm light. LOV2 and CatCh were stimulated at 470±24 nm light.

**Supplementary Figure 2.**
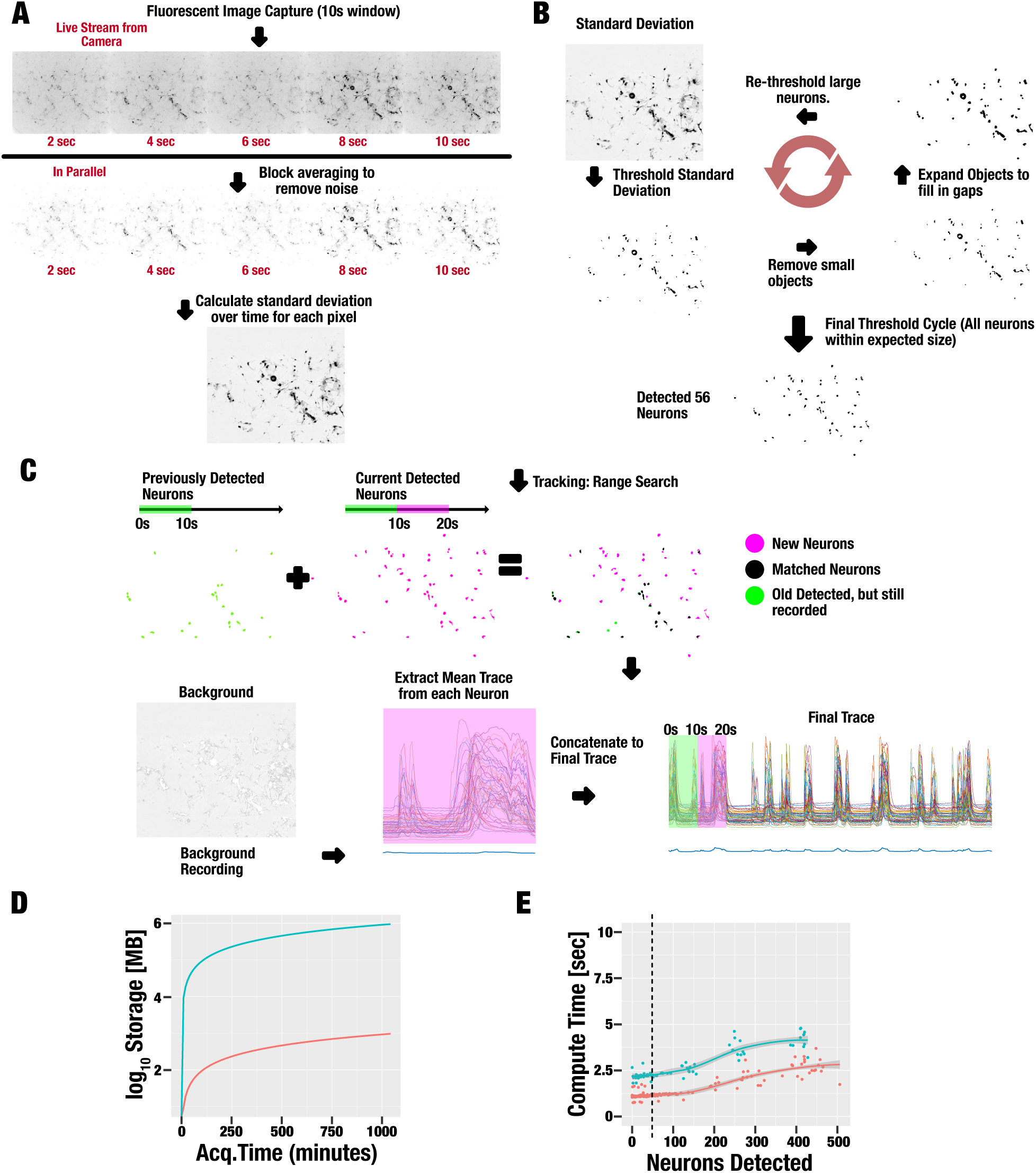
Illustration of neuron activity acquisition methods. **A.** Capture of Rcamp1.07 fluorescence and calculation of neuronal activity (expressed as standard deviation). **B.** Dynamic thresholding to segment individual neurons. **C.** Previously discovered neuron are tracked for the remainder of the acquisition and newly active neurons are added. Green regions are neurons detected in the first 10 seconds. Pink regions are neuron detected in the current data excerpt. These are aligned and pink regions that are not matched with previous regions are appended as new neurons. Calcium traces for all neurons and background are recorded and concatenated to the final trace. **D.** Storage requirements for raw video recording (teal) are 1,000x greater than our method (red). **E.** Benchmarking the number of neurons our system can process with 20 second excerpts (teal) and 10 second excerpts (red). Processing time must not exceed the batch time. Average number of neurons in a field of view is indicated by the dotted line.

**Supplementary Figure 3:**
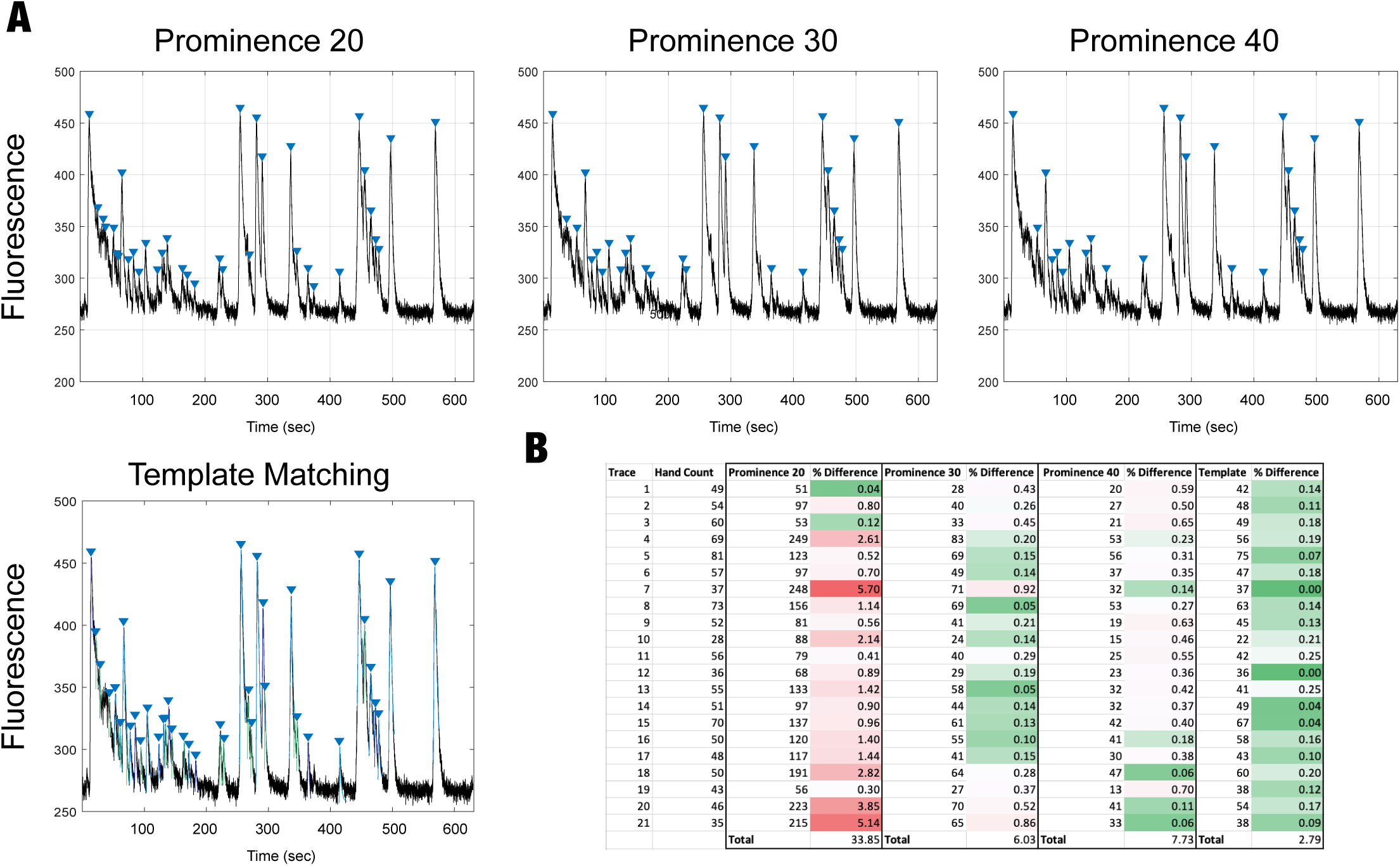
Comparison of calcium transient detection techniques. **A.** Analysis of an example trace from a control neuron with peak detection by prominence (units of prominence indicated) and with template matching. Identified calcium transient are indicated by a blue triangle directly above the peak. **B.** Comparison of techniques across 21 traces. Number of peaks detected were recorded and compared to the number of hand selected peaks (% difference). The technique with the lowest percent difference was template matching and was also the most consistent across traces. Colors indicate score on a scale of 0 to 5.14. All prominence levels were variable and worse performing than template matching.

**Supplementary Figure 4:**
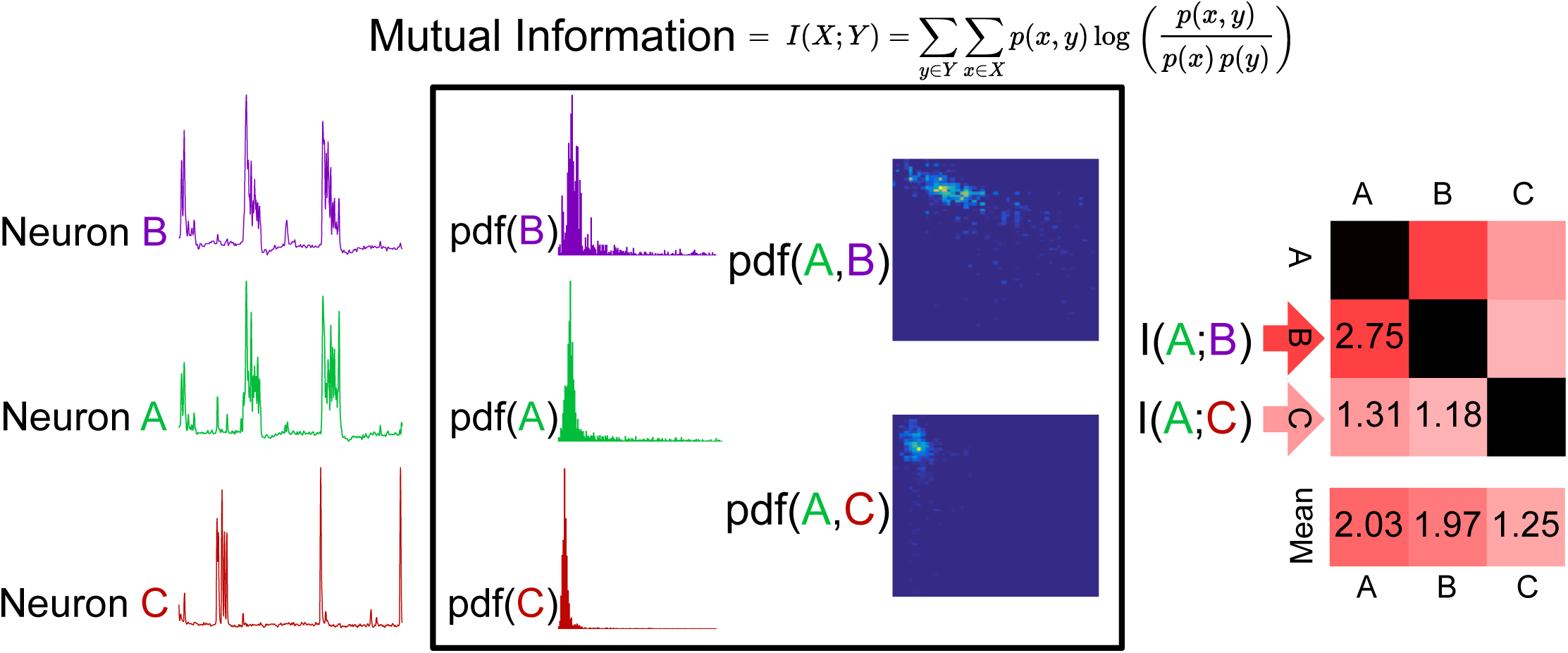
Calculation of mutual information (MI) across a field of view. Mutual information is calculated by pairwise comparisons of neuron traces. Each calculation uses probability density function to calculate a single MI statistic. These are then averaged across all permutated neuron comparisons to get an overall network connectivity for an individual neuron.

**Supplementary Figure 5:**
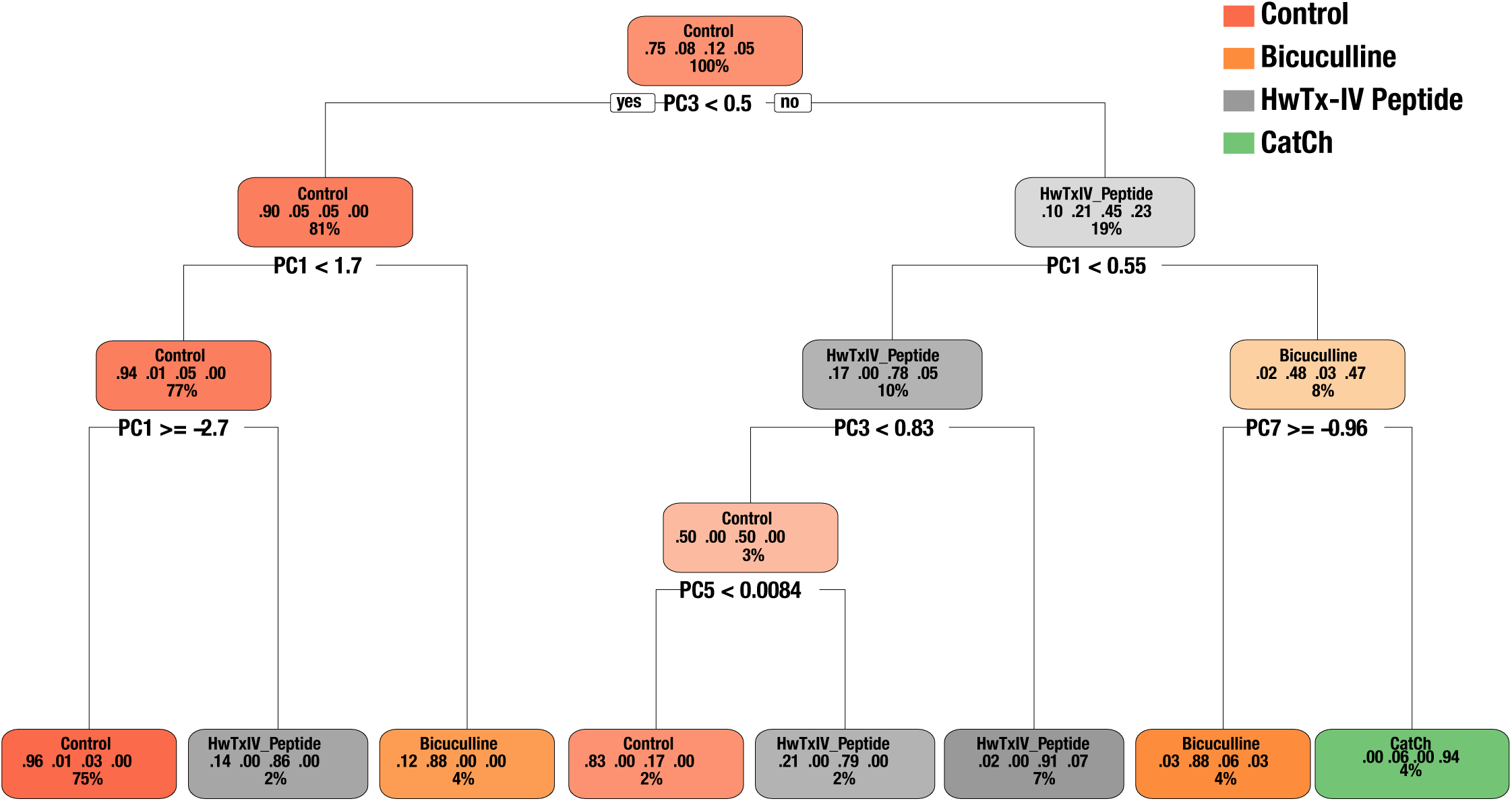
Decision Trees. Decision trees were trained on PCA data for calcium transient after the addition of the indicated peptide toxin or the during light stimulation (when CatCh was expressed). Tree were cross-validated 10-fold and restricted to a maximum depth of 4. All trees were restricted to a maximum depth of four. Decision tree leaves are described in the following way: the top-most number refers to the leaf class (peptide toxin addition or light stimulation for CatCh), next are percentage of permissive samples from Control, Bicuculline, HwTx-IV peptide, or CatCh within the leaf that fall within the class indicated in the top row. The bottom row indicates percentage of all data that are within the leaf. Color intensity of each leaf refers to purity of sample.

**Supplementary Figure 6:**
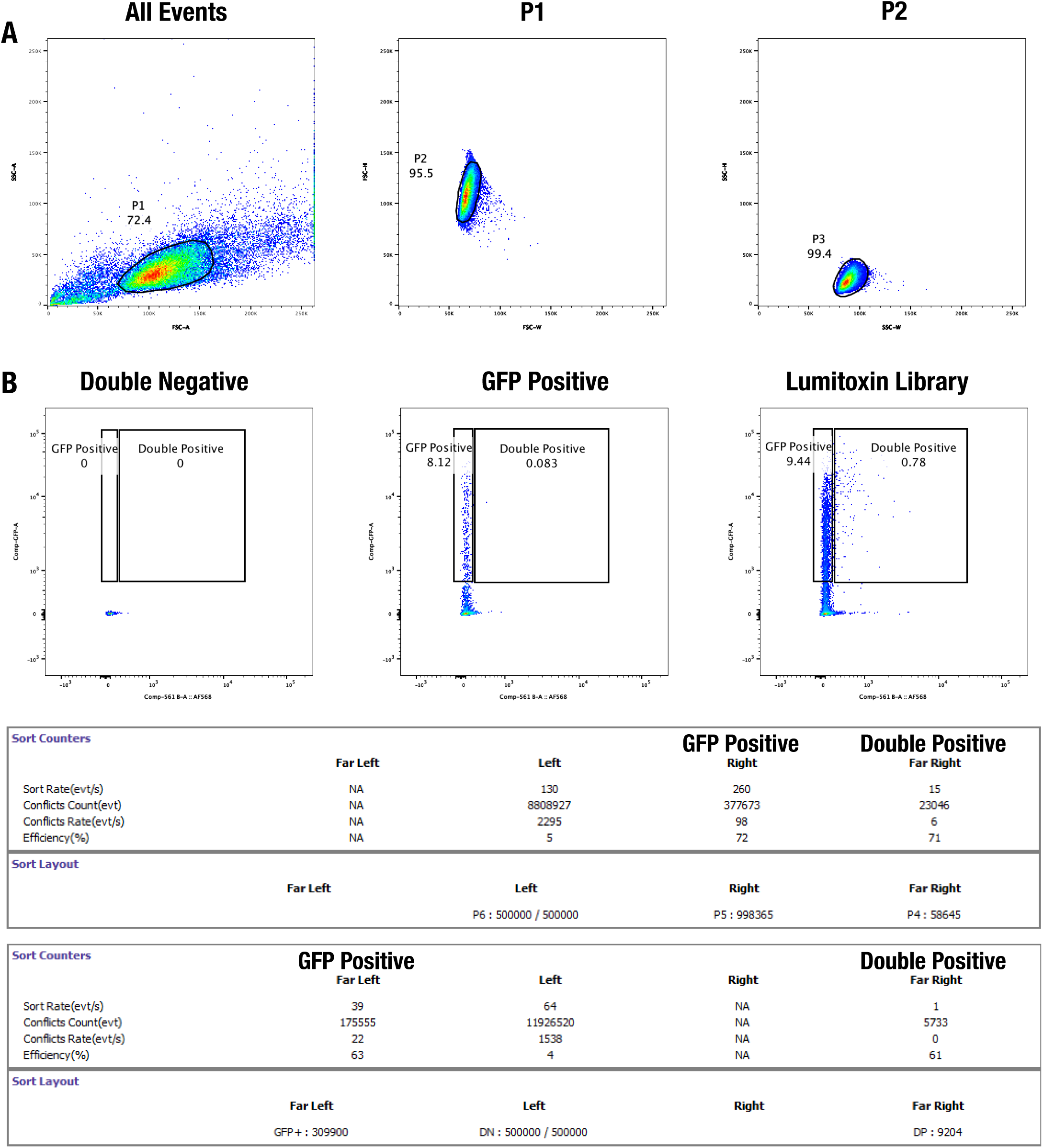
Fluorescence activated cell sorting of surface expressing lumitoxins. **A.** Gating scheme for isolation of single, viable HEK cells. **B.** The lumitoxin library was transiently expressed in HEK293 FT cells and labeled with mouse anti-FLAG (primary) and goat anti-mouse Alexa568 (secondary). Double negative and GFP positive cells were used as gating controls. **C.** Cells were sorted into two populations: GFP positive (internal expression) and Double positive (surface expression). DNA was recovered from these populations and sent for sequencing.

**Supplementary Figure 7:**
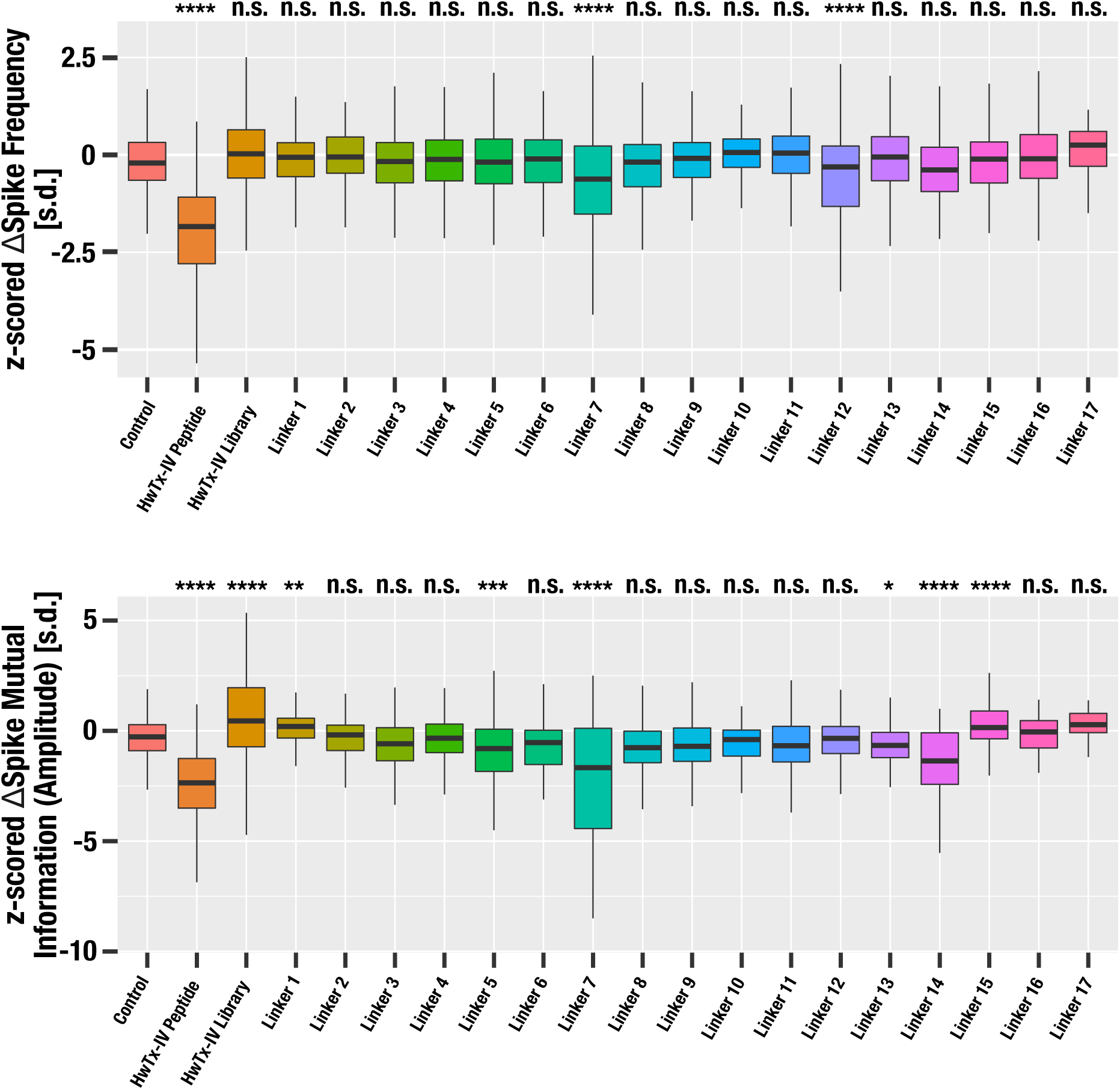
Light-induced changes in calcium transients properties depend on the lumitoxin linker. Select properties are shown as boxplots for controls neurons, neuron exposed to 20nM HwTx-IV peptide, neurons infected with the complete HwTx-lumitoxin library (includes all linker), or neurons infected with a lumitoxin comprised of HwTx-IV and the indicated linker. Box range represents interquartile range, median is indicated by a thick horizontal line, data range is indicated by a thin vertical line. Significance of the difference compared to control for a given class is indicated at the top of each panel (Dunnett’s many-to-one comparison with one control, **** p-value < 0.0001, *** p-value < 0.001, ** p-value < 0.01, *, p-value < 0.05, n.s. not significant).

**Supplementary Figure 8:**
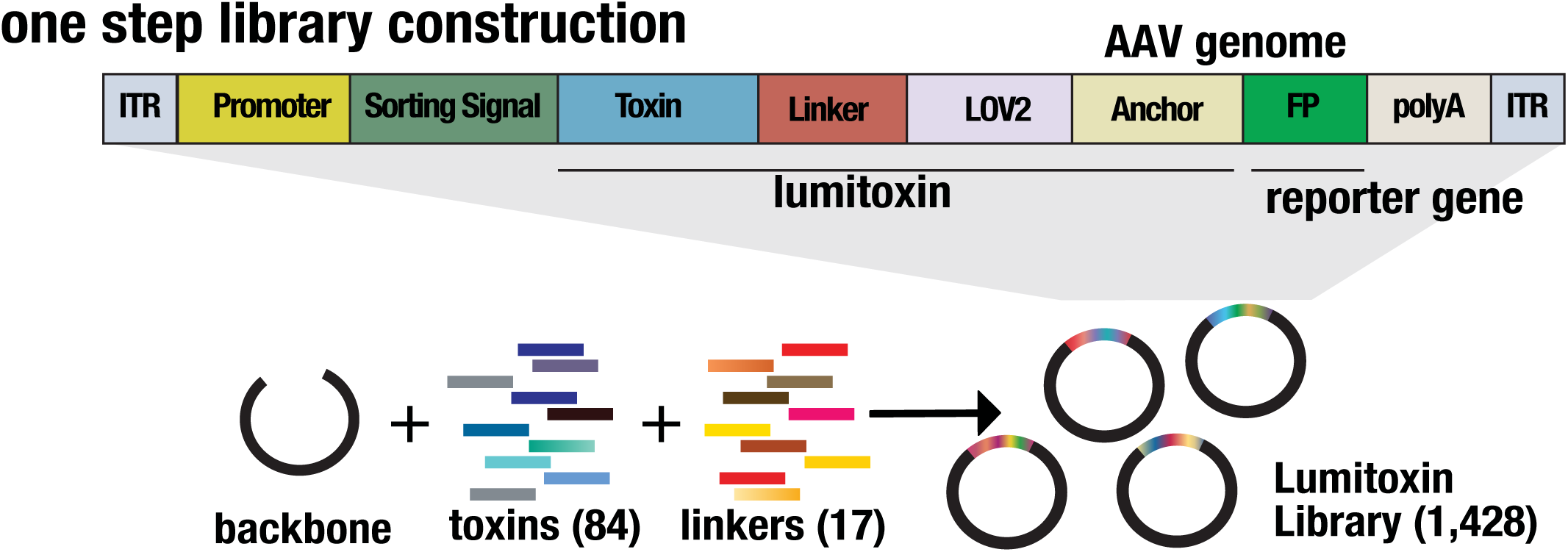
Lumitoxin library assembly. Libraries were assembled from synthesized gene fragments listed in Supplementary Tables 1 & 2. These fragments were sub-cloned into a shuttle vector (pUC19) with a unique BsaI restriction sites for toxins and a unique BsaI site for linkers. All fragments were assembled in a single reaction using Golden gate Assembly and recovered with over 30-fold coverage. The final library complexity was 1,428 members.

**Supplementary Table 1:**
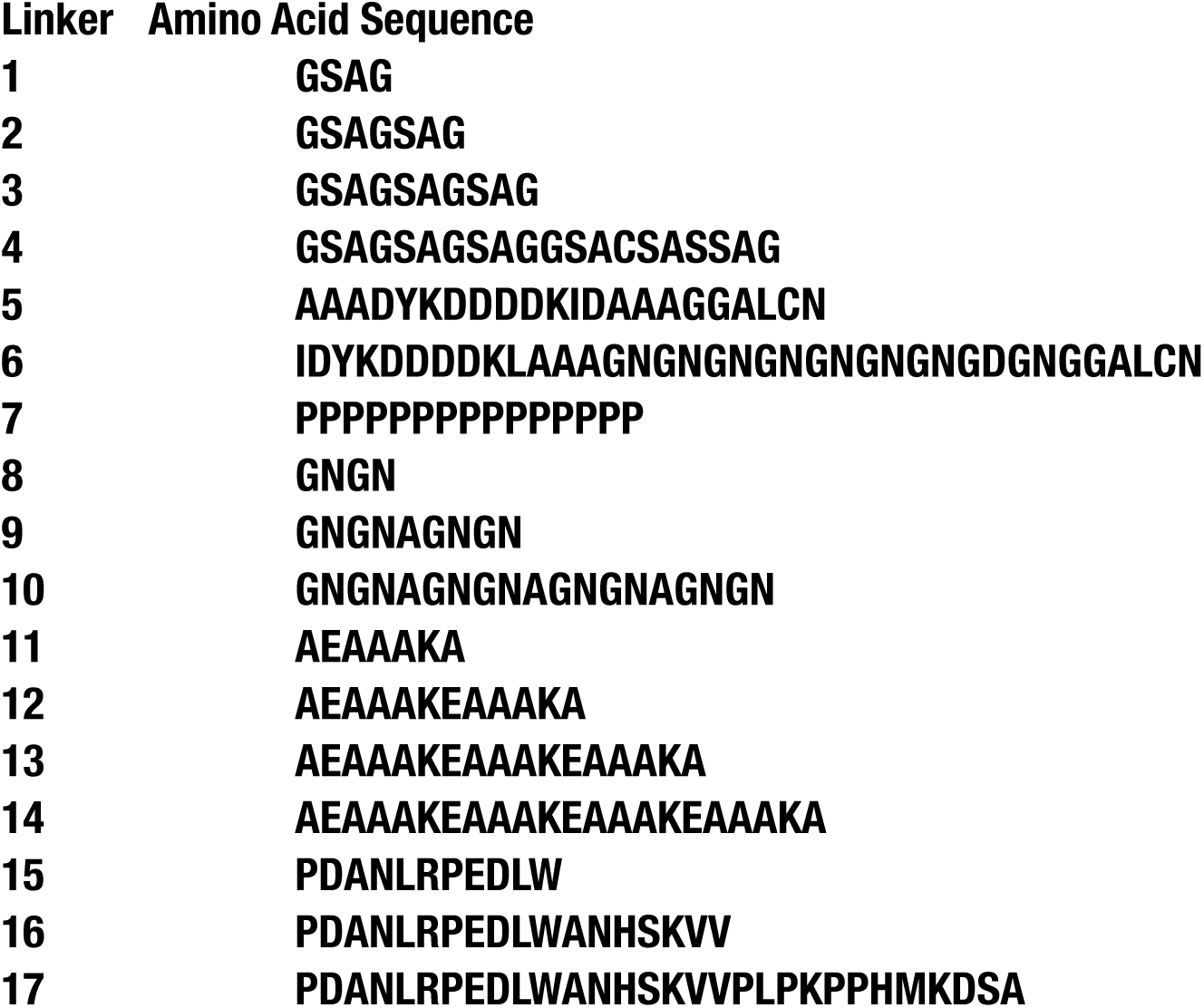
Amino Acid Sequence of lumitoxin linker.

**Supplementary Table 2:**
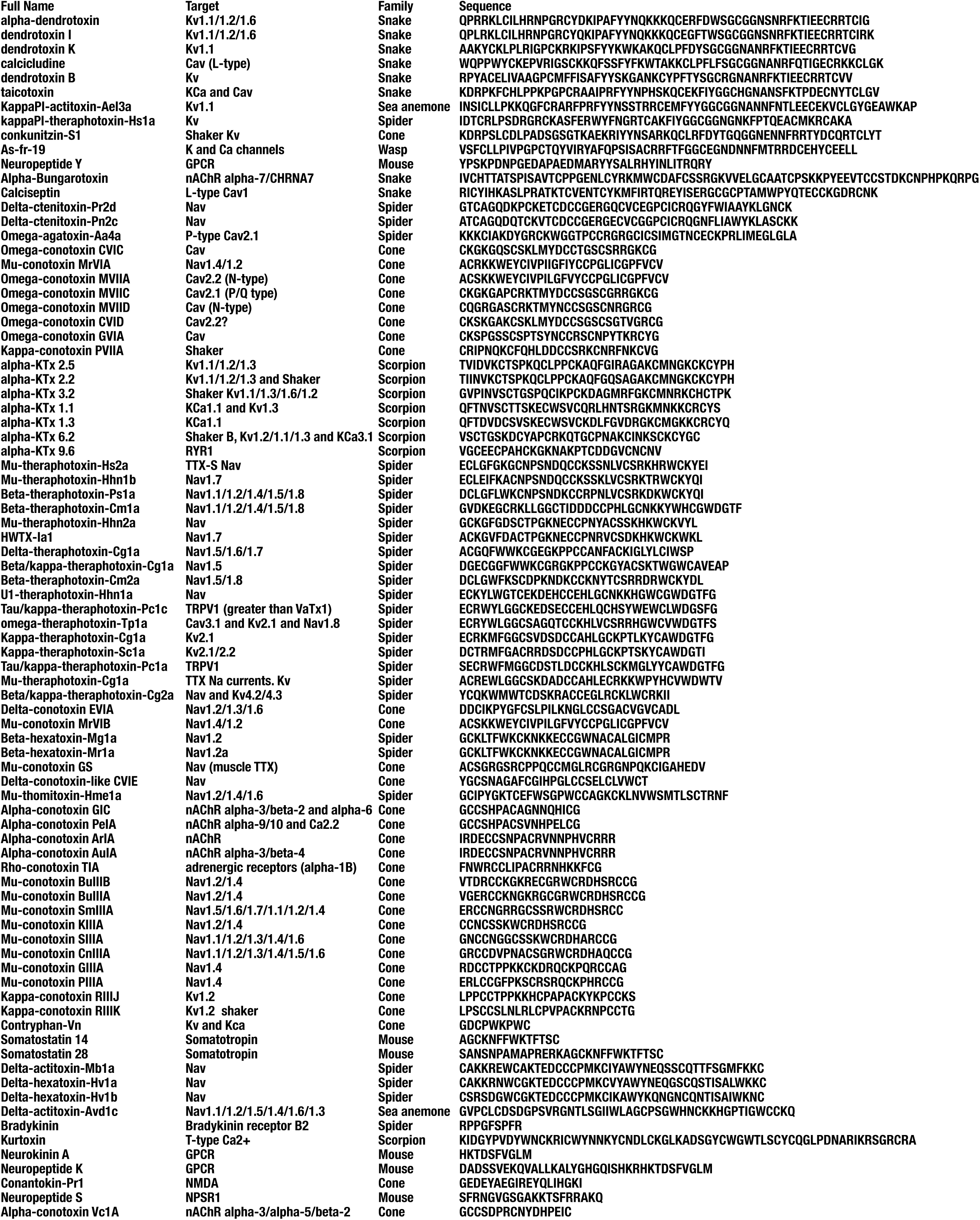
Amino Acid Sequence of encoded peptide toxins.

**Supplementary Table 3:** Phenotype prediction of neurons for the indicated tested condition.

| Condition tested | Predicted as |  |  |  |
| --- | --- | --- | --- | --- |
|  | Control | Bicuculline | HwTxIV | CatCh |
| Control | 28161 | 194 | 576 | 55 |
| HwTxIV Peptide | 652 | 66 | 3312 | 0 |
| HwTxIV Full Library | 33366 | 309 | 3005 | 0 |
| HwTxIV Linker 1 | 12863 | 85 | 0 | 0 |
| HwTxIV Linker 2 | 16547 | 40 | 223 | 0 |
| HwTxIV Linker 3 | 16066 | 0 | 346 | 0 |
| HwTxIV Linker 4 | 15648 | 0 | 256 | 0 |
| HwTxIV Linker 5 | 18212 | 126 | 546 | 0 |
| HwTxIV Linker 6 | 10169 | 205 | 203 | 0 |
| HwTxIV Linker 7 | 5650 | 143 | 734 | 0 |
| HwTxIV Linker 8 | 17213 | 137 | 1095 | 0 |
| HwTxIV Linker 9 | 11382 | 253 | 500 | 0 |
| HwTxIV Linker 10 | 16567 | 285 | 0 | 0 |
| HwTxIV Linker 11 | 20075 | 121 | 719 | 0 |
| HwTxIV Linker 12 | 23038 | 0 | 507 | 0 |
| HwTxIV Linker 13 | 20181 | 94 | 434 | 0 |
| HwTxIV Linker 14 | 2564 | 0 | 119 | 0 |
| HwTxIV Linker 15 | 5450 | 0 | 349 | 0 |
| HwTxIV Linker 16 | 1557 | 0 | 58 | 0 |
| HwTxIV Linker 17 | 2073 | 0 | 0 | 0 |

